# An Automated End-to-End Workflow for Production of Secreted Proteins in Transfected Mammalian Cells

**DOI:** 10.1101/2025.07.13.664612

**Authors:** Pavee Vasnarungruengkul, Michael A. Anaya, Annie W. Lam, Elisa Gonzalez, An Zhang, Maxine L. Wang, Woj M. Wojtowicz, Kai Zinn, Jost Vielmetter

## Abstract

The advancement of automation technologies has helped to enable a surge in large-scale screening efforts across fields such as molecular biology, protein biochemistry, cell biology, and structural biology. In the context of this “omics”-driven research, there is a need to generate automation platforms that are more flexible and less expensive, so that they can be utilized for basic research conducted by small groups. A key challenge in automation lies in developing methods that can replicate fine motor techniques that are normally performed manually by researchers at the bench. We are engaged in a large-scale project to map interactions among human cell-surface and secreted proteins and assess their effects on cells. This project involves production of a library of more than 2000 recombinant His-tagged fusion proteins secreted from transfected Expi293 cells. To execute such a project with a small group at an academic institution required construction of an affordable automated system that could also be used by other investigators. This led us to develop a high-throughput, 96-well format automation platform for end-to-end protein production. The workflow includes transformation of E. coli, plasmid DNA preparation, transient transfection, protein purification, desalting and buffer exchange, protein quantification, and normalization of protein concentrations, resulting in assay-ready proteins. The system is built around an in-house engineered modular robotic platform that integrates liquid handling with a suite of interchangeable ‘plug-and-play’ mobile enclosed device modules. Housed within a BSL-2 sterile environment, the platform enables flexible, fully automated workflows and can be readily customized for diverse user-defined protocols.

## Introduction

Interactions mediated by the extracellular domains (ECDs) of transmembrane receptor proteins control fundamental biological processes such as cell signaling, intercellular communication, and immune responses. Type I single-transmembrane (STM) cell-surface proteins (CSPs) have signal peptides (SPs) at their N-termini, followed by an ECD, transmembrane domain, and cytoplasmic domain. ECDs interact with secreted and cell-surface proteins expressed on neighboring cells and are targets for many antibody drugs approved for treating cancer, autoimmune disease, and other conditions.

For in vitro studies, CSP ECDs, as well as secreted proteins, are often produced as fusions with the Fc region of human IgG, which is a dimer that can stabilize proteins and increase solubility. As a result, dimeric ECD-human Fc (ECD-hFc) fusion proteins are often expressed by cells at higher levels than monomeric ECDs. The Fc region also facilitates efficient purification via Protein A or Protein G affinity chromatography. Additionally, the dimeric nature of the Fc region can increase avidity for binding partners, mimicking natural receptor states, and aids in capture and detection in binding experiments (Capon et al. 1989).

To meet the increasing demand for these and other proteins in high-throughput screening, automated production platforms have become essential. Automation enables a significant increase in the number of proteins that can be produced in parallel, substantially reduces manual handling and processing time, and accelerates the research timeline. It also improves reproducibility by minimizing human error and ensuring consistent processing conditions.

Additionally, miniaturized volumes can lead to considerable cost savings on reagents. Ultimately, automation empowers researchers to explore a wider range of experimental designs, leading to a more thorough investigation of biological systems and potential therapeutic targets.

The production of purified protein by transient transfection in mammalian cells involves purification of plasmid DNA, transfection, and protein purification, followed by desalting and buffer exchange, when necessary for experiments. Several groups have developed semi or fully automated workflows to carry out these methods. A custom-made robot for purifying transfection-grade plasmid DNA was first reported in 2006 (Kachel, Sindelar, and Grimm 2006) Since then, additional groups have developed semi-automated or fully automated plasmid DNA purification platforms (Cohen et al. 2022; Kates et al. 2023). In-house custom-designed robotic platforms for automated transfection (Grimm and Kachel 2002; Bos et al. 2015; Zhao et al. 2011) and protein purification platforms (Sun et al. 2012; Morimoto and Walinda 2024; Prince et al. 2025; Ransdell et al. 2023) have been engineered by several groups. There are also an increasing number of fully automated commercially available instruments that perform these individual steps.

We are conducting a large-scale project to map interactions among the ECDs of most human CSPs and secreted proteins and study the effect of these interactions on primary human mononuclear cells (PBMCs). To this end, we have developed a multiplexed protein-protein interaction screening method (Multiplex Interactome Assay; MPIA) that has the capacity to screen for interactions among every pairwise combination of thousands of proteins. To do this, we screen a single “prey” protein in solution against a multiplexed pool of 500 different “bait” proteins. The MPIA can examine binding between every pairwise combination of 384 preys with 500 baits within a few hours (i.e., 192,000 interactions screened). The MPIA is described in a companion paper that has been posted on bioRxiv.

To utilize the MPIA for a human cell-surface interactome screen, we need to transiently transfect Expi293 cells with >2000 expression plasmids, each containing sequences encoding an ECD fused at the N-terminus to a heterologous SP and at the C-terminus to human IgG Fc (hFc), followed by a series of epitope tags culminating in an 8xHis tag. These proteins (called ECD-hFc-8xHis) are secreted from Expi293 cells grown in suspension culture and are purified from the cell supernatant. To execute this work with a small group, automation is necessary.

Here we describe an in-house designed BSL-2 automation platform (Fig. 1) that produces purified proteins in a 96-well format. The end-to-end workflow is comprised of methods that transform bacteria with plasmid DNA, purify transfection-quality plasmid DNA from bacterial cultures, measure and normalize plasmid DNA concentration, transfect plasmid DNA into Expi293 cells, purify proteins from the cell supernatant via an 8xHis tag, desalt and buffer exchange purified proteins, and measure and normalize protein concentration. This workflow can be adapted to any tagged secreted protein produced by a cell line that grows in suspension.

**Figure 1.**
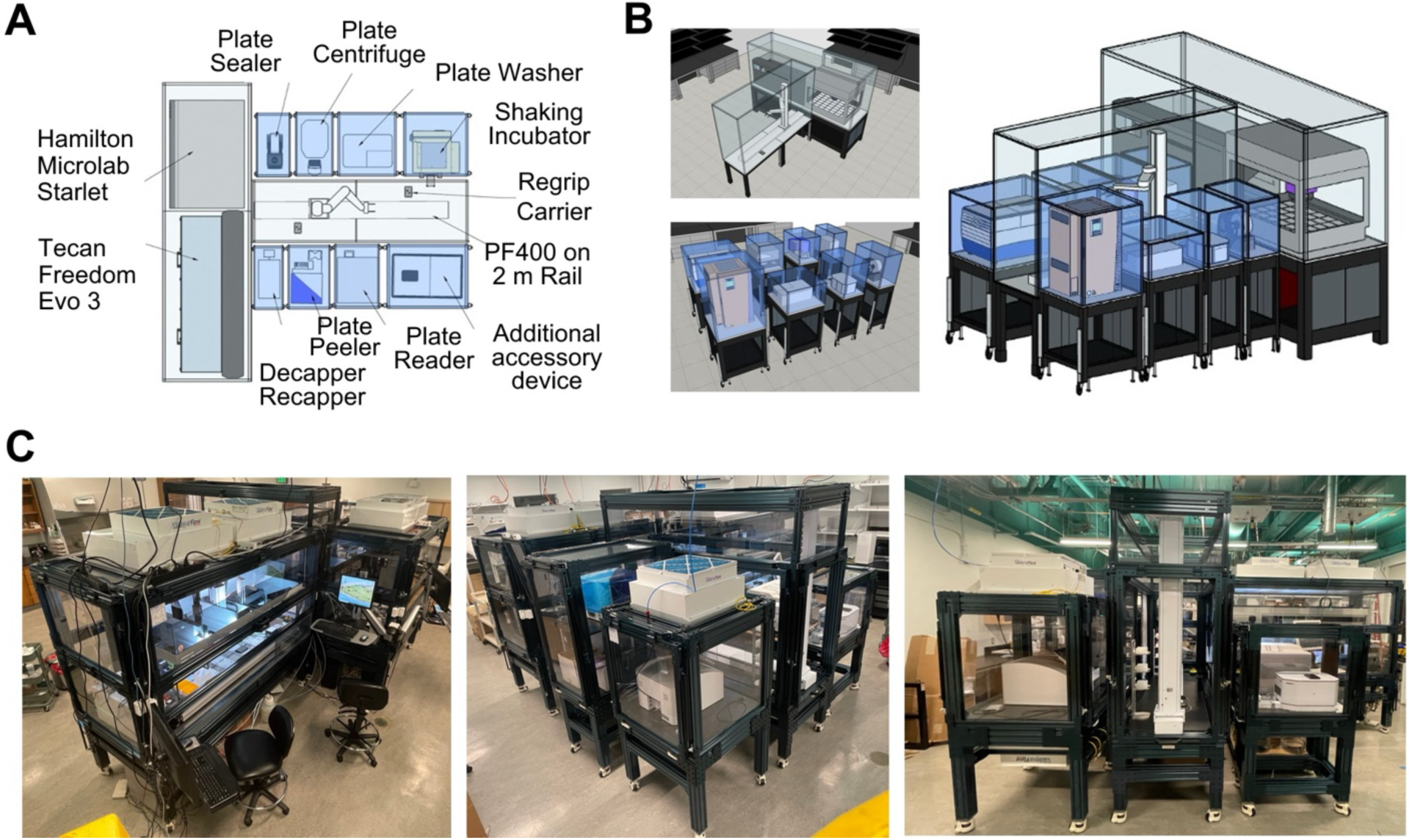
Automation platform. A. Schematic diagram. B. Schematic 3D views. C. Photographs of the assembled system.

The automation platform is modular. External instruments and devices, which are housed in individual BSL-2 enclosures, can be interchangeably airlocked to a central BSL-2 cabinet containing two liquid handlers. This is a ‘plug-and-play’ design, which allows user-defined flexibility as well as the option to add new instruments and devices to the automation platform (Wolf et al. 2024).

In the experiments described here, we transferred plates between modular devices by hand. However, we have incorporated a central robotic arm that can transfer plates between modules integrated into our platform, which will make it possible to generate fully automated workflows. With the fully automated system, we anticipate it will be possible to process four 96-well plasmid plates in parallel, resulting in the generation of 384 purified ECD-hFc-8xHis proteins per week. As such, the entire library of >2000 proteins for our cell-surface interactome screen could be produced in six weeks, thus allowing proteins to be tested for binding without prolonged storage at 4° C or freezing. Making proteins for this and other large-scale screens within a short time frame is important because long-term storage and/or freezing of purified protein can result in loss of binding or enzymatic activity.

## Results

### System Overview and Platform Architecture

Our workflow is built around a modular platform composed of liquid handlers, a plate reader, incubator, centrifuge, and other accessory devices (Fig. 1 and Table 1). All modules are arranged to facilitate fully automated manipulation of samples from DNA preparation to final protein purification, while maintaining operations within a BSL-2 sterile environment. Figure 1 provides both isometric and top view diagrams of the platform layout. The Hamilton Microlab Starlet and TECAN Freedom Evo 3 liquid handlers are housed in a stationary BSL-2 unit. Each accessory instrument has its own BSL-2 enclosure. The system is designed such that a PF400 robotic arm sits on a 2-meter rail in the main BSL-2 cabinet and is centrally located between the modular enclosed devices. The modular BSL-2 enclosures are designed to dock with the main PF400 enclosure using an in-house designed custom magnetic docking system. Through the magnetic docking tunnels, the PF400 can access each device, allowing for the seamless loading and unloading of plates and enabling transfer between any docked devices.

**Table 1.**
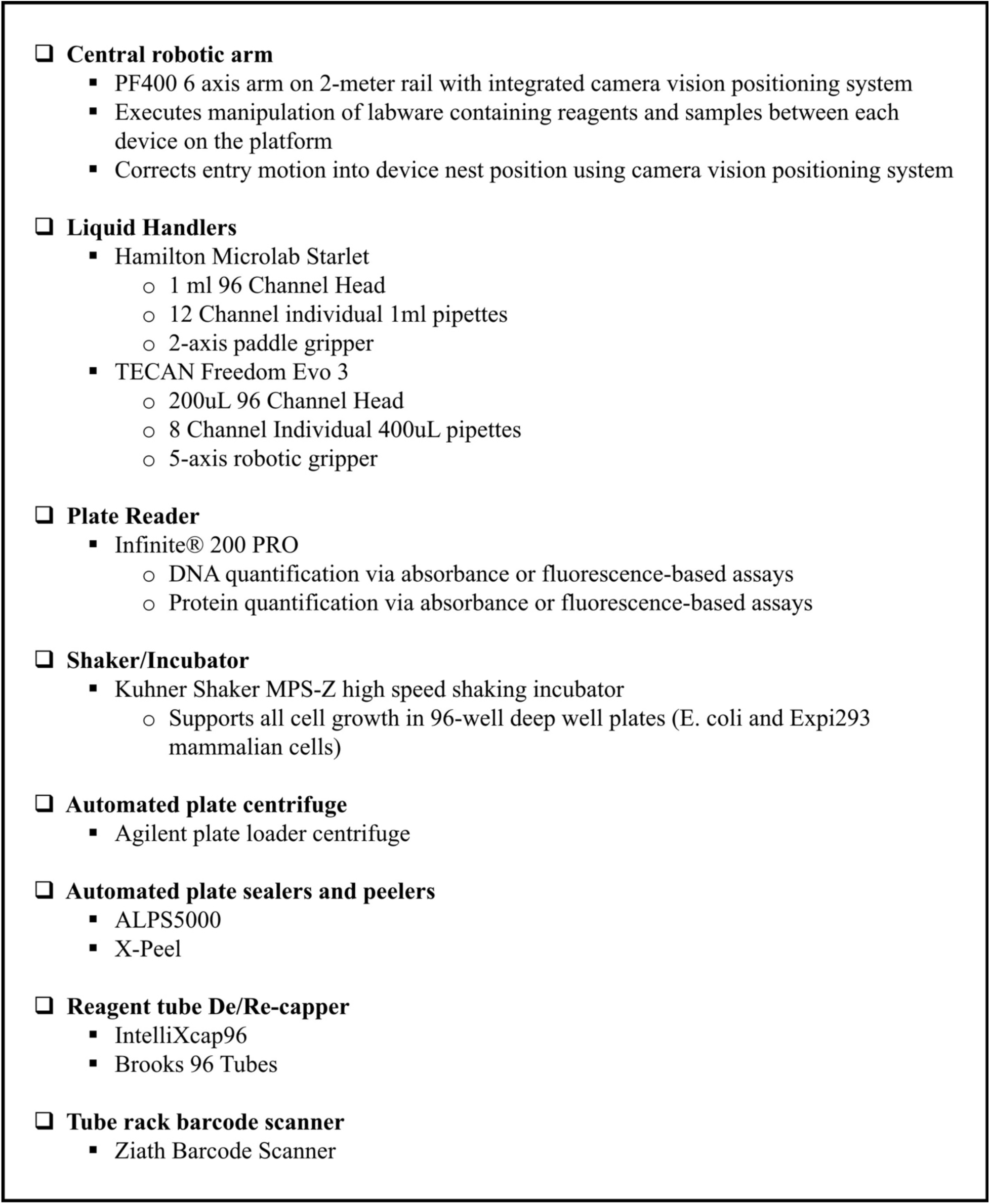
Automation devices on our platform.

The platform was engineered with modularity in mind, permitting individual devices to be detached for upgrades or repairs, thus supporting a flexible and upgradable system design. In addition, because the modular enclosed devices are on casters, devices can be quickly and easily swapped in and out depending upon the devices needed for a user’s individual workflow.

Modularity is space saving compared to systems that are built inside a single large enclosure with a fixed footprint, as is often found in the biotechnology industry. Additionally, it allows for devices purchased in the future to be integrated into the automation platform more flexibly. The Hamilton Starlet and TECAN Freedom Evo liquid handlers, while essential to the workflow, are docked in a similarly flexible way as the smaller devices and can also be replaced if future workflows should they require more specialized liquid handling modes.

The methods for the individual steps of the protein production workflow, which we describe below, were developed and optimized by manually moving plates between devices. For production of the >2000 protein library, we will integrate the PF400 robotic arm into the system, allowing the entire process to be fully automated. As such, in the workflow schematics in Figures 3-7, we have indicated the transfers that will be performed by the PF400 arm.

### Platform Workflow

The complete workflow described here is outlined in Figure 2. It is designed to purify secreted proteins bearing a His tag composed of 6 or more consecutive His residues. All methods are carried out in a 96-well format. The workflow begins with transformation and purification of plasmid DNA. Following plasmid purification, the concentration is measured and plasmids are diluted to the required concentration for transfection in Expi293 cells. Expi293 cells are transfected and secreted ECD-hFc-8xHis protein is allowed to collect in the supernatant for 4.5 days before harvesting. Proteins are purified from the supernatant via the 8xHis tag using Ni-NTA metal chelate affinity resin. Following purification, proteins are desalted and buffer exchanged, and protein concentration is determined using the Bradford assay and then normalized. The end product is a 96-well plate containing purified assay-ready proteins. For optimization of many steps in the workflow methods, we used the control protein mCherry-hFc-8xHis. mCherry-hFc-8xHis is a useful surrogate for ECD-hFc-8xHis proteins as a class, because its levels can be monitored using a fluorescence plate reader. All final products (glycerol stocks of bacterial cultures, purified plasmids, purified proteins) are stored in racks of individually-barcoded 96 screw cap tubes. Barcode scanning of racks ensures reliable sample tracking.

**Figure 2.**
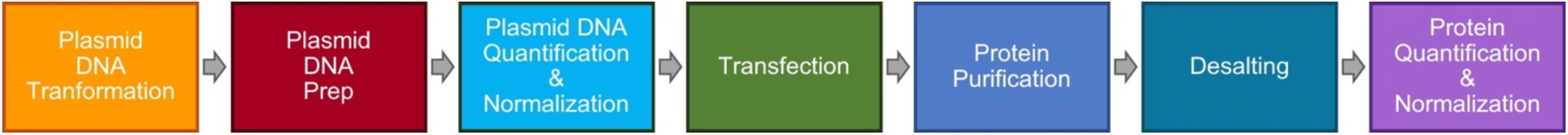
Overview of protein library production workflow.

Disposable plates are used for intermediate steps in workflows or when preparing reagents for subsequent workflows (e.g., normalized plasmid DNA for transfection is prepared in a 96-well PCR plate).

### Transformation of Plasmids and Generation of Glycerol Stocks

Automated plasmid preparation begins with transformation of E. coli cells (Fig. 3A). Competent DH5α cells are dispensed into 96-well PCR plates and transformation is performed by adding plasmid DNA to the cells, incubating on ice, adding medium, and incubating at 37 °C for 1 hour for recovery. Transformed cells are then plated in a 96-well format using the Tecan Evo liquid handler to ensure uniform colony distribution across the agar plate. Individual colonies are then hand-picked (future workflows may include a colony picker) and inoculated into 1 ml/well of selection media in 96-well deep well plates (96-DWPs) for overnight growth. Glycerol stocks of cultures are prepared and stored at –80° C as a renewable source. When needed, glycerol stocks are revived to inoculate fresh 96-DWPs and generate replacement glycerol stocks.

**Figure 3.**
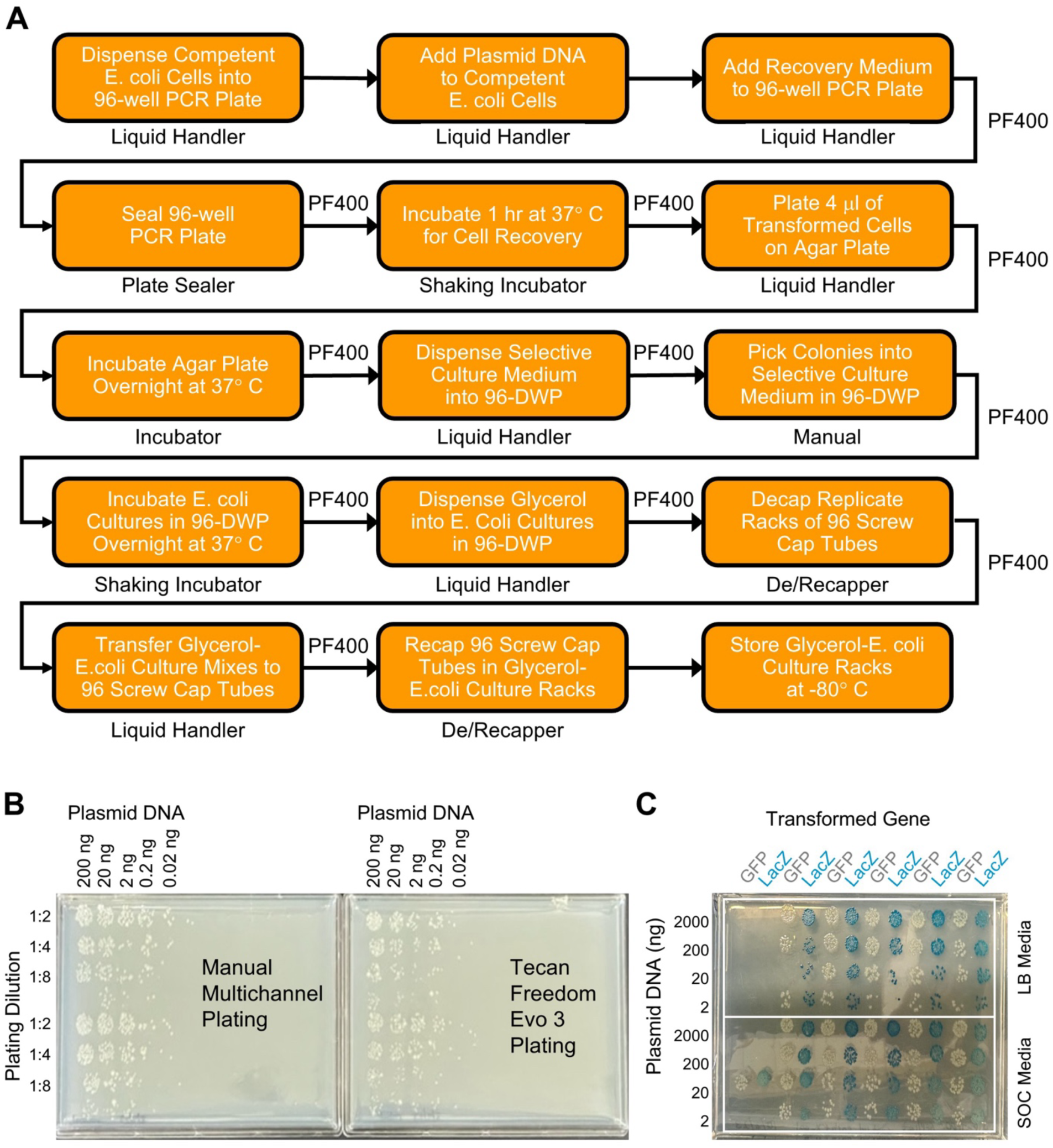
Automated transformation of E. coli. A. Workflow. B. Comparison of manual versus automated colony plating at different input plasmid DNA and dilutions of transformed bacterial culture. C. Bacteria transformed with plasmids expressing GFP (white) and LacZ (blue) were plated in alternating columns to visualize cross-contamination. Two recovery media (LB and SOC) were compared at various plasmid DNA input amounts.

We found that transformation efficiency was greatly affected by the method used for dispensing the DH5α competent cells into 96-well PCR plates. Following production of competent cells, we compared two methods. In the first method, we dispensed cells into 96-well PCR plates and stored the plates at –80 °C. In the second method, we froze cells in 5 ml aliquots, stored the tubes at –80 °C and, following thawing, dispensed cells into a 96-well PCR plate. Transformation efficiency was much higher when the cells were frozen in 5 ml aliquots and dispensed into the 96-well PCR plates after thawing (data not shown).

The majority of steps in this method, and in many of the methods described below, involve liquid handling aspiration and dispense operations. Optimization of these steps involved fine tuning parameters on the liquid handlers such as tip selection, aspiration and dispense speeds, tip height, and use of liquid sensing. Additional considerations included custom liquid class definitions, prewetting, blowout settings, tip travel speeds, and surface tension compensation to maintain sample integrity and ensure accurate volume transfer. Techniques such as reverse pipetting, tip-touch, liquid tracking, and controlled tip positioning within wells were used to further improve precision.

In this and other methods, some steps required the development of novel liquid handling strategies to replicate manual techniques performed by researchers. The transformation workflow involves plating transformed bacterial cultures on an agar plate. When performed manually, plating is commonly done on large 10 cm agar plates and the bacterial culture is spread on the plate using a glass “hockey stick” or glass beads. For higher throughput, some researchers have adopted the use of “mini agar plates” (agar poured into in 6-well, 12-well or 24-well plates). Our plasmid libraries are in racks of 96 screw cap tubes. As such, we sought to develop an automation method that would allow us to carry out the entire transformation method in a 96-well plate (including plating of the bacterial culture using the Tecan Evo 96-tip MCA head).

Automated micro plating in a 96-well format requires dispensing a small volume of bacterial culture with precise control of tip height above the agar surface. The tips must be positioned high enough to avoid piercing the agar, yet close enough for the hanging droplet to make contact and wick onto the surface. To generate a 96-well compatible agar plate, we first tried filling a 96-well gridded plate with agar. This method failed to produce a level agar surface due to interference from the grid structure. We found that pouring agar into a non-gridded rectangular plate (i.e., single reservoir with the same dimensions as a standard 96-well plate) resulted in a level agar surface that allowed plating a small volume of transformed bacterial culture. Importantly, agar plates needed to be prepared fresh as an evaporation “edge effect” occurred during storage at 4° C, even when plates were stored at 4° C for only 3 days.

To optimize a method for plating, we first used a multichannel pipet and manually dispensed transformed bacterial culture onto the agar plate surface. We tested a range of plating volumes (2-10 µl) and found that plating 4 μl of bacterial culture allowed sufficient spreading on the agar surface, while ensuring that the plated cultures did not spread into one another, which was essential to prevent cross-contamination. To optimize cell density for single colony formation, we performed transformation using a 10-fold dilution series of miniprepped plasmid DNA (0.02-200 ng) and plated a dilution series (Fig. 3B). We found that plating 4 μl of undiluted culture following transformation with 10-20 ng of plasmid DNA gave rise to individual colonies with sufficient spacing to allow picking a single colony. Similar results were observed when plating was performed using the 96-tip MCA head on the Tecan Evo. To evaluate cross-contamination between plated cultures, we performed one transformation with a plasmid expressing GFP and a separate transformation with a plasmid expressing LacZ. These two transformed cultures were plated using the Tecan Evo in alternating columns on an agar plate containing X-gal (Fig. 3C). When grown on an agar plate containing X-gal, colonies expressing LacZ are blue while colonies expressing GFP are white. We observed no cross contamination of blue and white colonies. Additionally, colony counts were similar when either LB media or SOC media were used during the 1 hour recovery incubation prior to plating.

### Plasmid DNA Preparation

Automated plasmid DNA preparation is performed using the Macherey-Nagel NucleoBond 96 Xtra EF, 96-well kit. The workflow is shown in Figure 4A. The protocol is fully automated on the Hamilton Starlet liquid handling system and all pipetting steps, incubation timings, and reagent transfers are encoded in the Instrument Method software.

**Figure 4.**
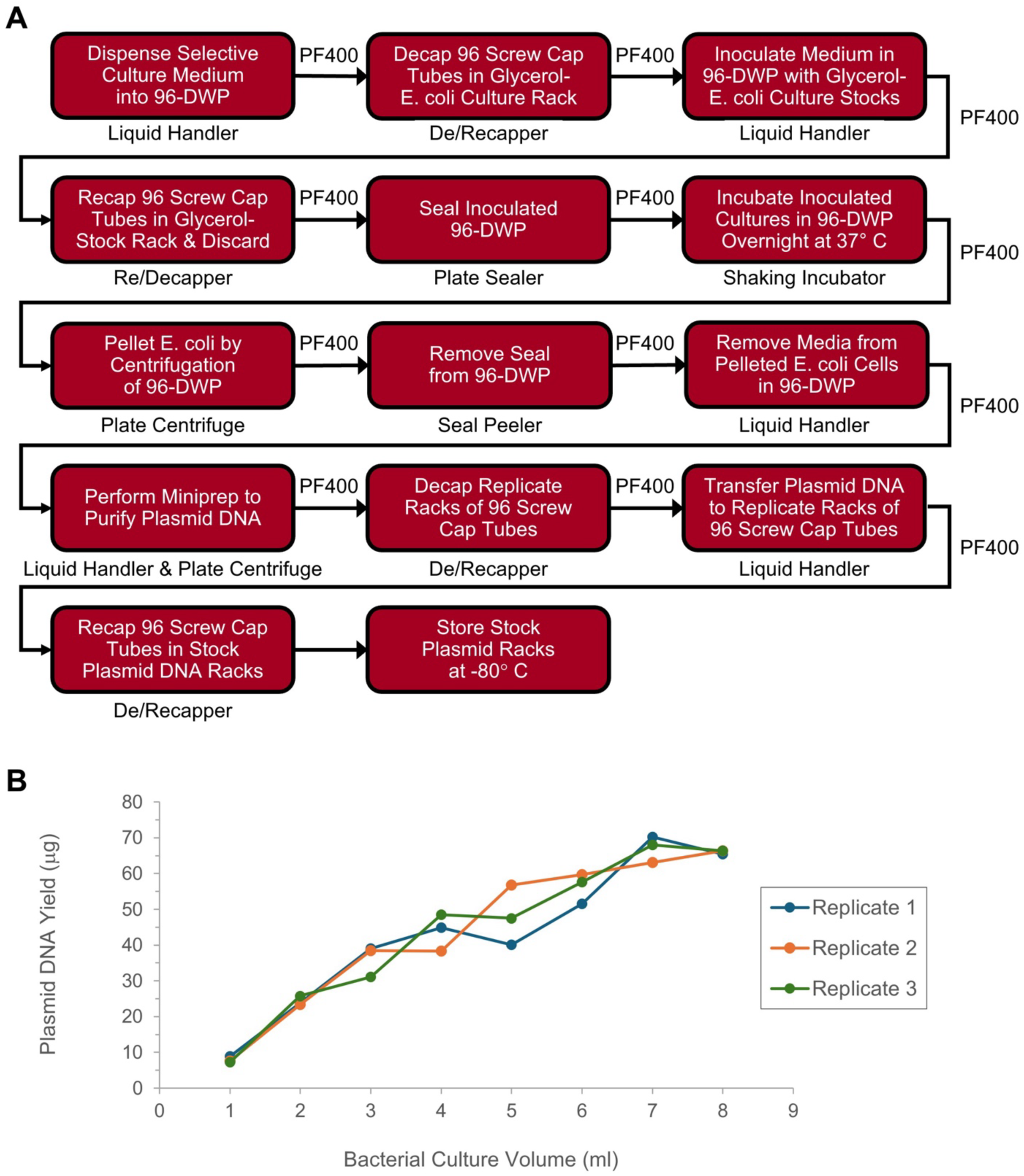
Automated plasmid DNA preparation. A. Workflow. B. Correlation of increasing input of E. coli culture with plasmid DNA yield.

Overnight bacterial cultures (1 ml/well) are grown in four replicate 96-DWPs (see below). Bacteria are pelleted by centrifugation, resuspended in resuspension buffer, and the four replicate plates are pooled into a single 96-DWP plate. Cell lysis is induced by addition of lysis buffer, and gentle pipetting with 1 ml pipette tips. The lysed cell suspension is then neutralized by adding neutralization buffer and mixing, and the lysate is clarified by vacuum filtration through a vacuum filter plate. Plasmid DNA is then transferred to a silica-based spin column plate, washed under gravitational flow conditions, spun dry and eluted with elution buffer. The eluted DNA is precipitated by adding isopropanol and transferred to the finalizer plate for further purification of the plasmid DNA. After washing the precipitated plasmid DNA, it is redissolved and eluted with sterile water into a clean 96-well elution plate and transferred to 96 screw cap tubes to prevent evaporation during storage.

As the 96-DWPs used for growing the bacteria cultures accommodate 1 ml of culture per well and the NucleoBond 96 Xtra EF, 96-well kit is capable of purifying up to at least 50 ug of plasmid DNA per well (a yield far greater than the amount of plasmid in a 1 ml culture), we reasoned that cultures could be grown in replicate 96-DWPs, and the lysate filtered re-using a single lysate filtration plate several times. To determine the number of replicate culture plates that could be used, we performed plasmid DNA purification from increasing volumes of bacterial culture (1-8 ml) and measured the plasmid yield (Fig. 4B). With increasing volume of culture, the yield increased linearly up to a volume of 4 ml. It continued to increase up to 8 ml. However, at volumes greater than 4 ml, the rate of increase in yield diminished, indicating a nonlinear, saturating trend. Furthermore, the risk of clogging the lysate filtration filter plate after too many reuses increases. Based on these results, we use four replicate 96-DWPs of bacterial culture in our workflow.

Following pelleting of the bacteria by centrifugation, it is necessary to fully resuspend the cell pellets. When performed manually in a single tube, a researcher can angle the tube, position the pipette tip just above the cell pellet along the tube wall, and gently pipette up and down – adjusting speed, angle, and tip position as needed – to gradually resuspend the cells without dislodging the pellet. This technique is important because, once dislodged, it is difficult to completely dissociate the floating cell pellet. As such, it was necessary to develop a liquid handling method that could mimic this manual technique. Using x- and y-axis tip positioning around the edge of the well, combined with optimized aspiration and dispense speeds, we developed a technique on the liquid handler that allows resuspension of cells without dislodging the pellet. To determine the completeness of resuspension, we evaluated the presence of particulates by visual inspection.

Another step in the protocol that required development of an automation technique to emulate manual benchwork is processing of the sample following addition of lysis buffer. When performed manually at the bench, the tube is immediately capped after lysis buffer is added and the tube is gently inverted several times until the lysate is homogeneous. During this step, the lysate mixture becomes highly viscous and gel-like due to the release of intracellular contents, including genomic DNA. We developed a liquid handling method that mimics manual tube inversion using 1 ml wide bore tips. The wide bore is essential to prevent clogging and it reduces shearing of genomic DNA. Additionally, it allows the lysate mixture to be gently mixed with minimal delay following addition of lysis buffer, while also maximizing volume turnover with each aspiration and dispense step during mixing.

### Plasmid DNA Quantification and Normalization

Plasmid DNA concentrations are determined using absorbance measurements at 260 nm and 280 nm to derive O.D. 260 values and 260/280 ratios. This workflow is shown in Figure 5A. Following plasmid DNA elution into 96-well PCR plates, a small volume is transferred into a UV-transparent 96-well assay plate and diluted with TE buffer 1 to 10. Absorbance readings are performed to calculate concentration and purity. Typically, the concentration of purified plasmid DNA ranges from ∼200-1000 ng/μl, which translates into a yield of ∼14-70 μg (Fig. 5B). For unknown reasons, we sometimes observe wells with significantly lower yields (e.g., wells E08 and F08 in the representative plate results shown in Fig. 5B). However, with rare exceptions, the concentration of plasmid DNA in these low yield wells is above the minimum concentration required for preparation of transfection complexes (100 ng/μl). We know that low yield is not the result of specific ECD transgenes because, when plasmid preps for these are repeated, we obtain standard yields.

**Figure 5.**
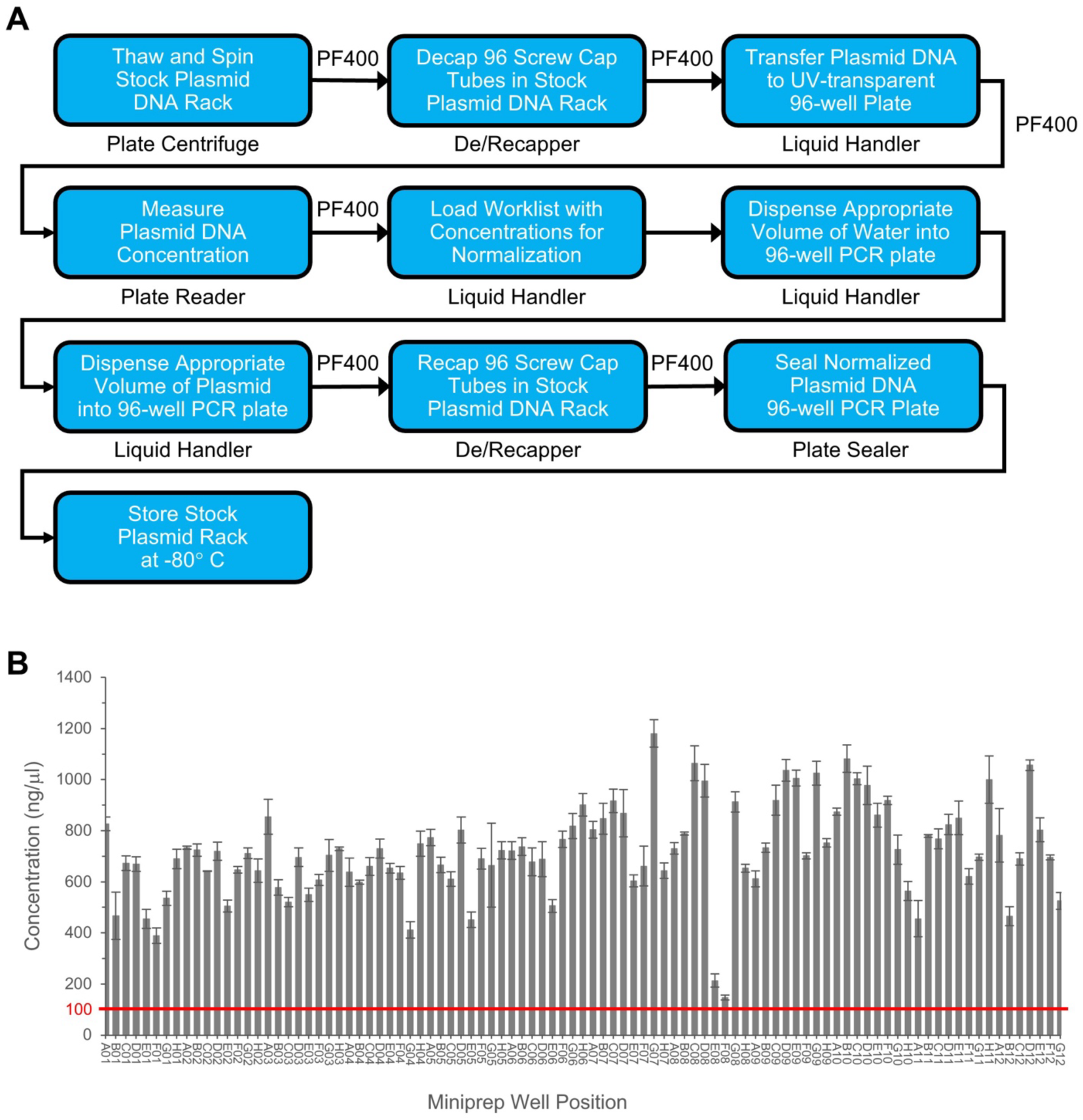
Plasmid DNA quantification and normalization. A. Workflow. B. Plasmid DNA yield data from a representative library plate. The red line at 100 ng/μl indicates the minimum concentration required for preparation of transfection complexes.

Concentration normalization of plasmid DNA is carried out using Opti-MEM as the diluent. A workbook is generated that maps the location and concentration of each stock plasmid in the rack of 96 screw cap tubes. Normalization scripts are executed on the Tecan Freedom Evo to calculate and dispense the appropriate volumes of plasmid DNA and diluent per well into a target PCR plate, ensuring uniform concentrations across all samples for downstream transfection. Appropriate volumes are transferred to obtain a final concentration of 20 ng/µl in a total volume of 50 µl for one transfection plate. If the aim is to generate 4 transfection plates the volume has to be scaled to 200 µl total.

### Mammalian Cell Transfection

Automated transfection of Expi293 cells is carried out in 96-DWPs (1 ml/well) and cells are cultured under standard conditions (37 °C, 8% CO₂, 1000 rpm shaking, 3 mm orbital radius). Cells are seeded at a density of 3 × 10^5^ cells/ml. Four replicate plates of cells are seeded for transfection of each 96-well plate of plasmids, resulting in a total transfection volume of 4 ml of cells for each plasmid. Transfections are carried out using the ExpiFectamine 293 Transfection Kit. Plasmid DNA-lipid complexes are prepared by diluting the plasmid DNA and ExpiFectamine reagent in Opti-MEM, followed by a 3 minute incubation. After complex formation for an additional 5 minutes, mixtures are automatically distributed to the four replicate cell plates and plates are transferred to the incubator. Enhancers are added 17–20 hours post transfection and plates are returned to the incubator. Supernatant is harvested 4.5 days post transfection at which time the cell viability is ∼70%. This workflow is shown in Figure 6A.

**Figure 6.**
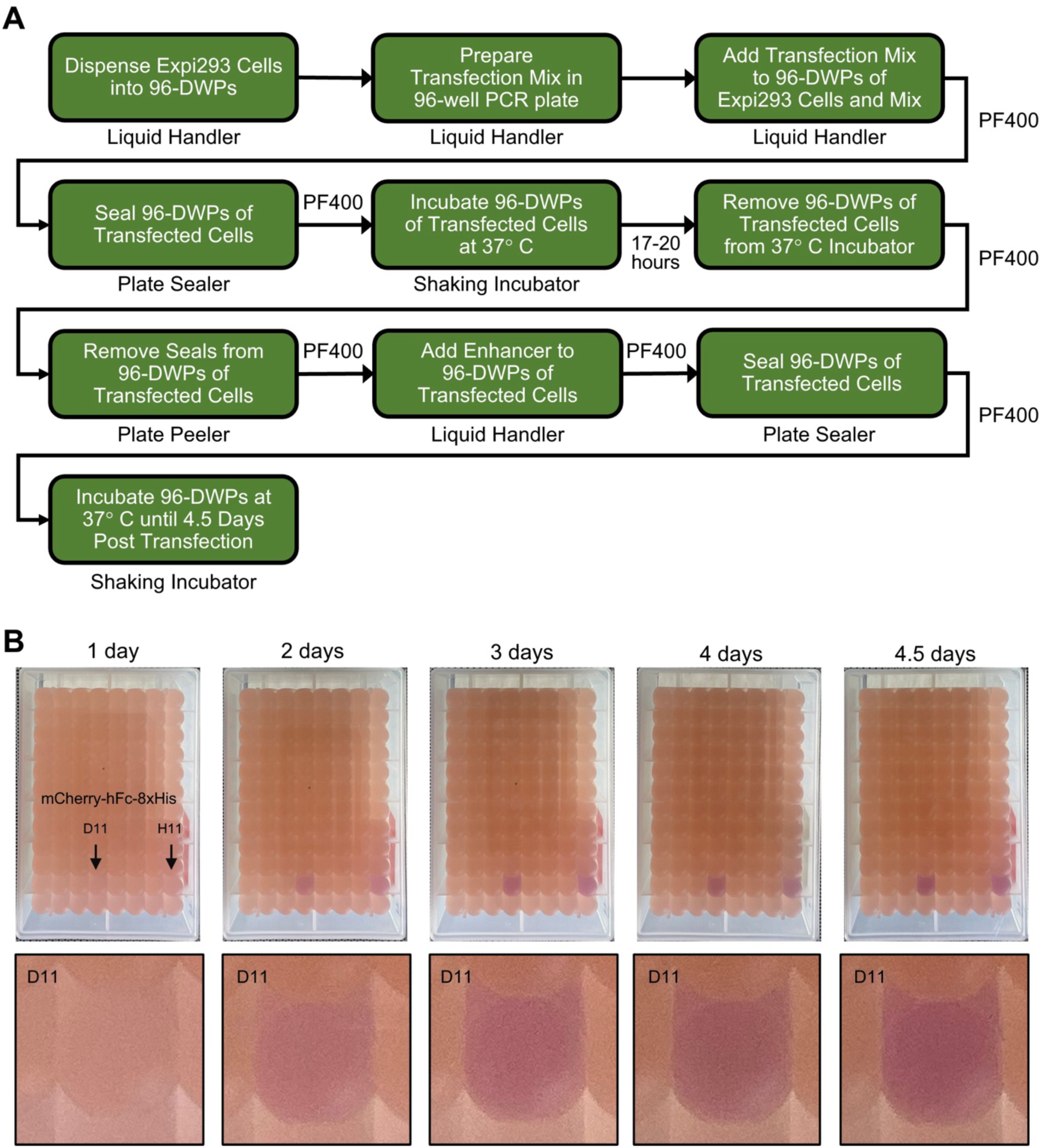
Automated mammalian cell transfection. A. Workflow. B. Images of transfected 96-DWP from 1 day to 4.5 days post transfection, showing expression of mCherry-hFc-8xHis protein at different times after transfection. A control plasmid expressing mCherry-hFc-8xHis was transfected in two wells (arrows). Zoomed in view of well D11 is shown below each plate image.

To monitor protein expression over time, we transfected a plasmid encoding the secreted protein mCherry-hFc-8xHis. As mCherry-hFc-8xHis is secreted and accumulates in the cell culture media, the color of the media changes from light pink to magenta, which is visible by eye (Fig. 6B). mCherry-hFc-8xHis expression continued to increase up to 4.5 days post transfection. Beyond this point, cell viability declined significantly. Therefore, in our workflow, we harvest the cell supernatant at 4.5 days post transfection.

During development of the transfection workflow, we discovered that the fluid dynamics created by 3 mm orbital shaking in the square wells of the 96-DWP are critical for maintaining cell health. At a 3 mm orbital radius, the circular orbit of the shaking platform is not visible when the platform is viewed from the side, so deviation from the expected shaking pattern is not readily observable. However, when a strict circular orbit is maintained, which is essential for optimum cell viability, the location of a dot marked at the center of a well appears stationary during shaking when observed from above in time-lapse video. In one of our experiments, the viability of the cells was poor, approaching 20% on day four post transfection and, although the plate was seated properly on the shaker platform, in some wells the cells were no longer in suspension. Upon investigation, we discovered that the shaker, when loaded with only two plates diagonally across from each other, had an elliptical rather than circular shaking pattern. It is thus critical to maintain the circular shaking pattern. We found that a shaker has to be robust to generate a consistent circular 3 mm shaking orbit, and that shaker performance degrades over time. Even new shakers, when fully loaded with four plates, can have a reduced orbit diameter, caused by counter-rotation movements of the shaker body induced by the momentum of the platform load. We were able to significantly reduce this effect by tying down the shaker body to the incubator shelf with zip-ties. We have now obtained a Kuhner high speed shaking incubator (Table 1), and are integrating it into our platform. The experiments described here were done with orbital shakers placed inside an incubator. We anticipate that there will be fewer problems with the Kuhner shaker/incubator system. Unlike our current incubators, this shaking incubator can be loaded using the robot arm, as indicated in the workflow of Figure 6A, and it is required to create a completely automated system.

### Cell Harvesting and Protein Purification

Automated protein purification begins with harvesting of the cell supernatant 4.5 days post transfection. Protein is then purified from the supernatant using Ni-NTA metal chelate affinity resin. This workflow is shown in Figure 7A. The four replicate 96-DWPs are removed from the incubator and cells are pelleted by centrifugation. Importantly, removal of the supernatant is performed using the Hamilton Microlab Starlet, which is equipped with tip-based aspiration and liquid sensing and tracking to avoid disturbing the pelleted cells. Supernatant is transferred from the four 96-DWPs into four 96-well PALL filter plates (1.2 µm) and filtration is carried out by centrifugation into four fresh 96-DWPs. Due to evaporation during the 4.5 day post-transfection incubation and the need to leave a small volume behind to avoid disturbing the cell pellet, following filtration, the amount of clarified supernatant in each well of the 96-DWPs is approximately 800 µl.

**Figure 7.**
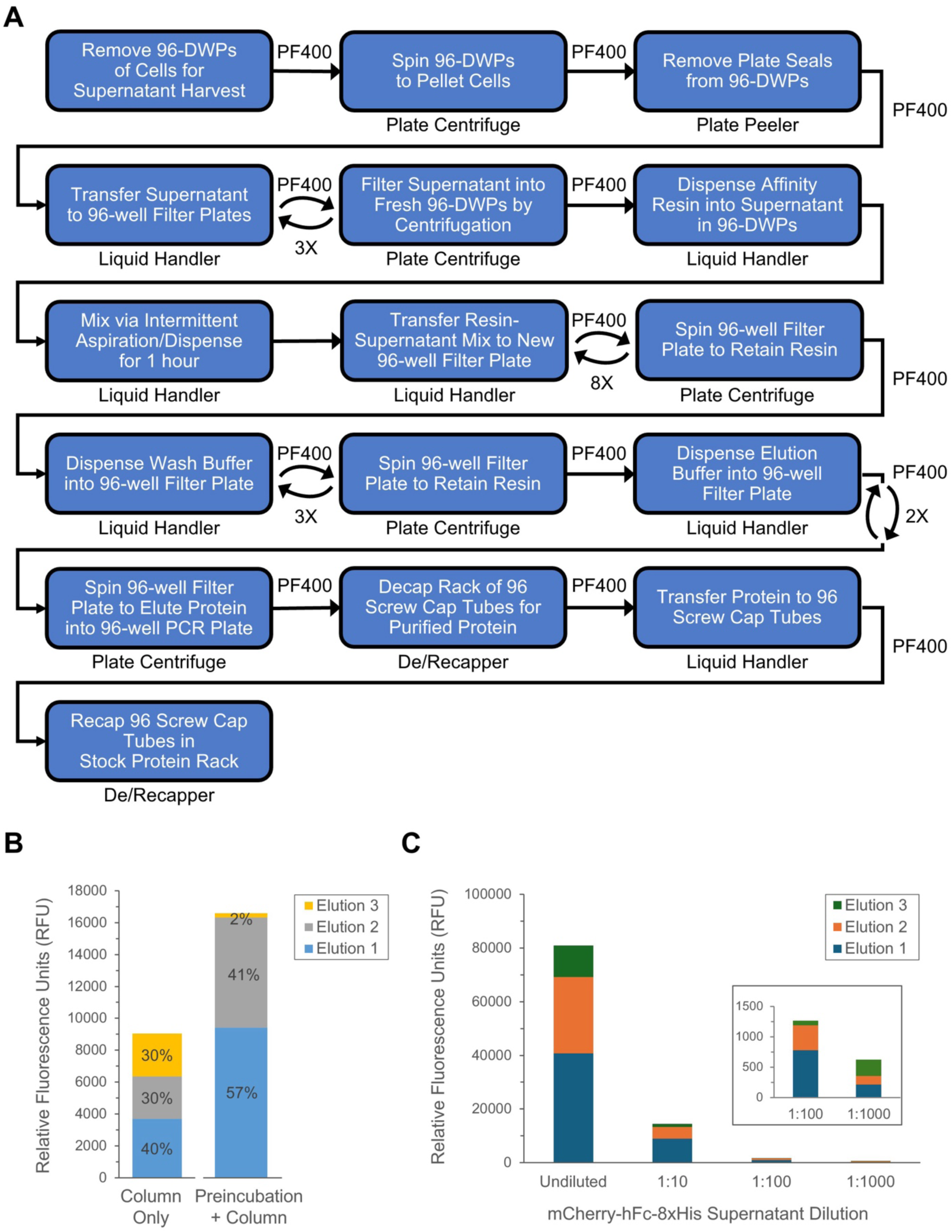
Automated supernatant harvesting and protein purification. A. Workflow. B. Fluorescence of purified mCherry-hFc-8xHis following each of three elutions with 500 mM imidazole. C. Purification with different dilutions of mCherry-hFc-8xHis.

The protein purification method continues in the four fresh 96-DWPs of clarified supernatant. We now need to pass the clarified supernatant through Ni-NTA metal chelate affinity resin, which binds to the 8xHis tag at the C-terminus of each ECD-hFc-8xHis protein, to purify the proteins. Based on the binding capacity of Ni-NTA resin, we calculated that 50 ul of resin should be more than sufficient to bind the amount of ECD-hFc-8xHis present in 3.2 ml of supernatant (the total volume of four replicate plates each containing 800 µl of supernatant). However, when we tested increasing bed volumes in the filter plate well, we observed that 200 µl of resin was required to retain an evenly distributed column bed.

We then tested a purification method in which clarified supernatant was passed through the resin column bed in the filter plate without preincubation of supernatant with resin. We sequentially centrifuged 400 µl of supernatant from cells expressing mCherry-hFc-8xHis through a 200 µl resin bed in the filter plate, collecting the flowthrough in a fresh plate after each spin and measuring mCherry-hFc-8xHis fluorescence using a plate reader. Although the binding capacity of the resin far exceeded the amount of mCherry-hFc-8xHis present in the supernatant, we observed high levels of mCherry signal in the flowthrough. This suggested that the mCherry-hFc-8xHis may not have had sufficient time to bind the resin as it passed rapidly through the bed during centrifugation.

These observations led us to develop a combined preincubation+column method which maximized protein yields and was feasible to implement on the robot. We structured our method so that we would build the final resin bed volume up to 200 µl in the filter plate by successive addition of supernatant/resin transfers. In this method, 100 µl of equilibrated resin (50% slurry; 50 µl bed volume) is added to the 800 µl of clarified supernatant in each well of the 96-DWPs. Resin is incubated with the supernatant at room temperature for 1 hour with intermittent resin resuspension achieved by pipetting up and down every 5 minutes on the liquid handler.

Following incubation, the resin–supernatant mixture from the four replicate plates is sequentially transferred to a 96-well PALL filter plate (1.2 um pore size). This is done by transferring 400 µl volumes (two per well containing 800 µl supernatant), with centrifugation performed between transfers to allow the supernatant to pass through the filter. This process pools the 200 µl of resin from the four replicate plates into a single filter plate and packs it into a resin bed.

We compared the column bed method to the preincubation+column method using undiluted mCherry-hFc-8xHis supernatant, eluting the protein bound to the resin with 3 x 100 µl volumes of 500 mM imidazole. Each elution was collected in a fresh plate and mCherry-hFc-8xHis fluorescence was measured using the plate reader. We observed a ∼2-fold increase in purified mCherry-hFc-8xHis protein using the preincubation+column method (Fig. 7B), so we selected this as our method.

Using the preincubation+column method, we observed that almost all of the mCherry-hFc-8xHis protein was eluted by the first two volumes. However, we were unsure if this would also be the case for less abundant proteins. Optimizing purification efficiency (and thus protein yield) for low-abundance proteins in the supernatant was a key priority, as low-expressing proteins may fall below our target concentration for downstream protein-protein interaction and cellular assays. We used dilutions of mCherry-hFc-8xHis supernatant to determine the optimal volume of elution buffer needed to elute low-abundance proteins from the Ni-NTA resin (Fig. 7C). mCherry-hFc-8xHis in undiluted supernatant (∼100 ng/µl) and three ten-fold dilutions of supernatant (1:10, 1:100 and 1:1000) were incubated with resin and transferred to the filter plate as described above. We chose 1:1000 as our highest dilution because proteins expressed at levels below this concentration (∼0.1 ng/µl) cannot be quantified using the Bradford assay (see below). We performed three sequential elutions with 100 μl of 500 mM imidazole, each time collecting the eluted protein in a separate PCR plate. mCherry fluorescence measurements showed that, under undiluted conditions (i.e., highest starting concentration), most of the protein was recovered in the first two elutions. At low mCherry concentrations in the supernatant (i.e., 1:100 and 1:1000 dilutions), protein elution was more evenly distributed across all three elution steps. This shift toward less efficient elution for low concentration samples suggested that recovery of low concentration proteins would be more complete if we used multiple elutions and larger elution volumes. Based on these results, we chose to elute proteins with a more generous volume of 400 μl of 500 mM imidazole, using 2 x 200 μl elutions for higher efficiency (rather than 4 x 100 μl elutions).

### Desalting and Buffer Exchange

Automated desalting and buffer exchange is performed when removal of imidazole is necessary for downstream applications, as it is toxic to some primary cells in culture, including PBMCs. Desalting is performed using spin column plates based on size-exclusion filtration. To desalt proteins, buffer exchange to PBS is performed using 96-well 40K molecular weight cut off (MWCO) ThermoFisher Zeba plates. Desalting plates are pre-equilibrated by washing with PBS three times, using centrifugation to remove the buffer between each wash. After washing, 100 μl of protein is added to each well and the plate is centrifuged to collect the eluate in a clean 96-well plate. This workflow is shown in Figure 8A.

**Figure 8.**
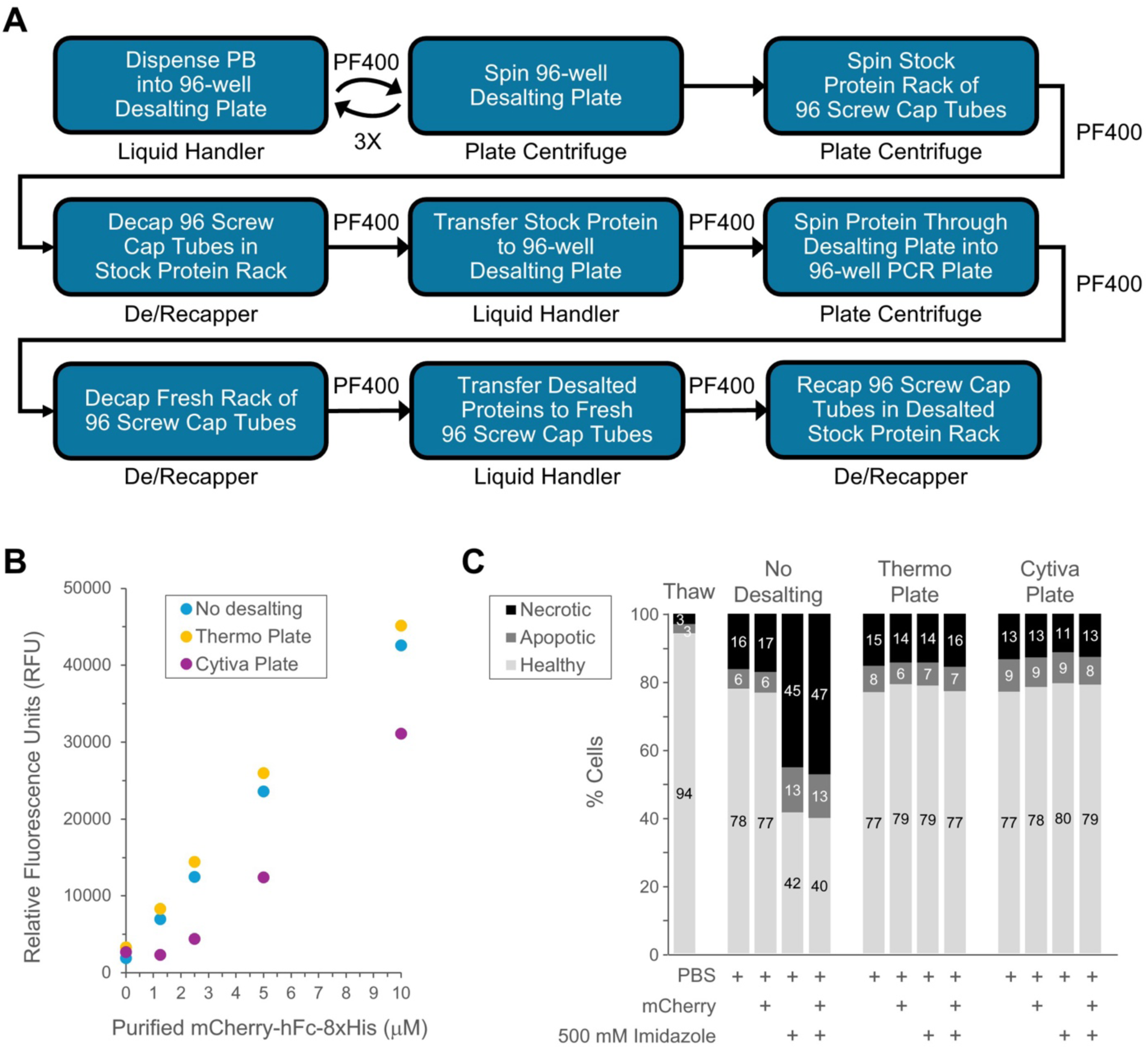
Automated desalting and buffer exchange. A. Workflow. B. Recovery of purified mCherry-hFc-8xHis using 40 kDa MWCO Zeba (ThermoFisher) and 5 kDa MWCO PD MultiTrap G-25 (Cytiva) 96-well desalting plates. C. Effect of desalting mCherry-hFc-8xHis on viability and health of PBMCs after a 48 hr. incubation period.

We selected the 96-well 40 kDa MWCO Zeba (ThermoFisher) desalting plate based on a comparison with the 96-well 5 kDa MWCO PD MultiTrap G-25 (Cytiva) desalting plate. In side-by-side comparison experiments, 2-fold dilutions of purified mCherry-hFc-8xHis ranging from 10 μM to 1.25 μM were desalted using the Zeba and PD MultiTrap G-25 plates, according to the manufacturer’s protocol, and recovery was determined using mCherry fluorescence in a plate reader (Fig. 8B). No protein loss was observed using the Zeba plate. Although the PD MultiTrap G-25 plate has a lower MWCO, significant protein loss was observed. Protein loss was inversely related to input concentration, with 67% loss observed at 1.25 μM and 27% at 10 μM.

In the absence of desalting and buffer exchange, proteins containing imidazole exhibit toxicity toward PBMCs, as evidenced by increased cell death and apoptosis. To assess desalting efficiency, mCherry-hFc-8xHis protein processed using the Zeba and PD MultiTrap G-25 desalting plates was incubated with freshly thawed PBMCs. Cell necrosis and overall health were assessed 48 hours post addition via flow cytometry using dyes specific for viability and apoptosis markers. Figure 8C shows the results for proteins containing imidazole that were incubated with cells, both pre and post desalting. Cell viability and health were comparable to PBS controls for proteins processed using both the ThermoFisher Zeba and Cytiva PD MultiTrap G-25 plates.

### Protein Quantification and Normalization

Proteins are quantitated using the Bradford assay. The workflow is shown in Figure 9A. Four 4-fold dilutions of each stock protein are prepared in triplicate 384-well plates. A dilution series of purified BSA is used to generate a standard curve. Bradford reagent is added to each well and absorbance is measured at 595 nm. Protein concentration is calculated using the BSA standard curve.

**Figure 9.**
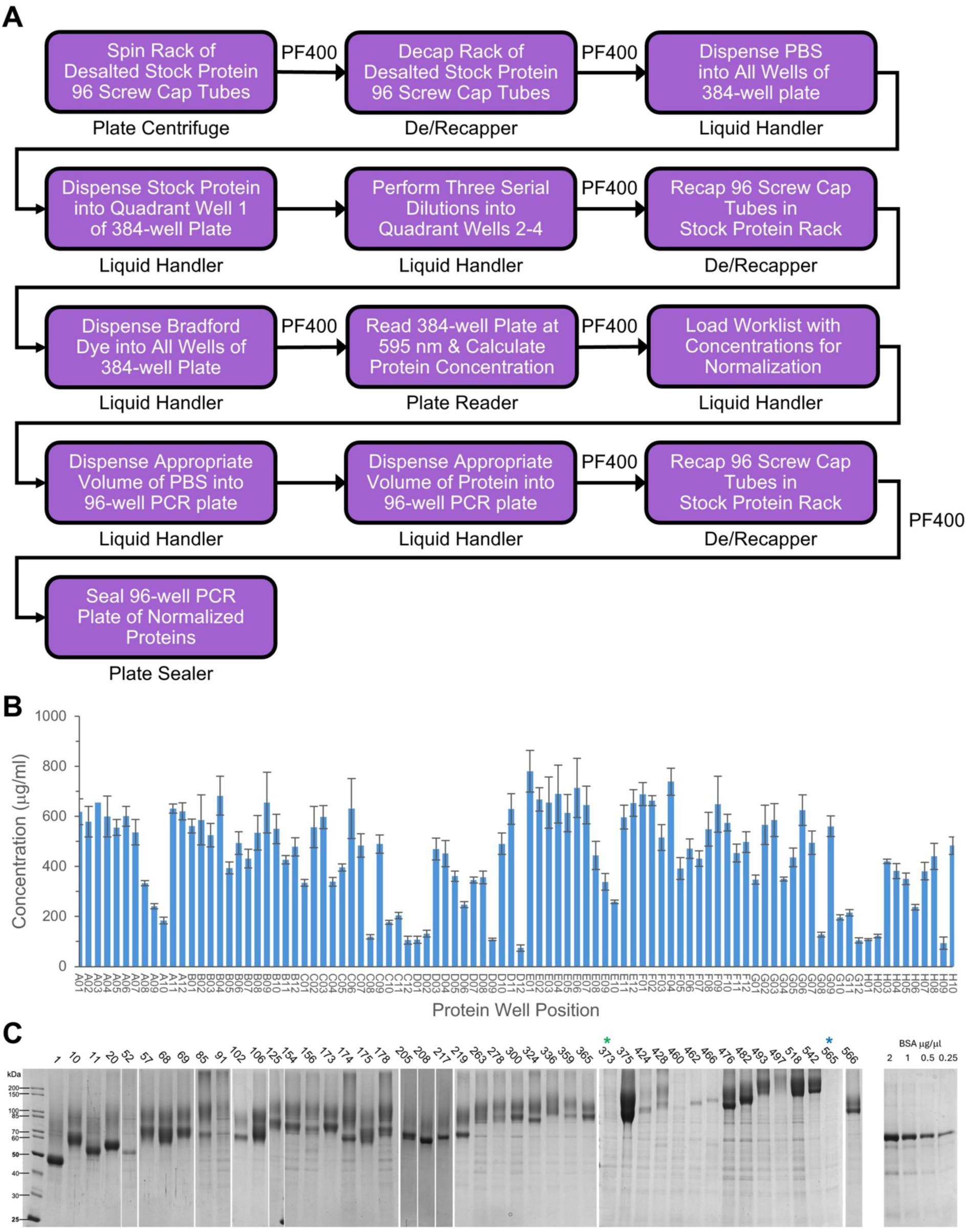
Workflow for protein quantification and normalization, and analysis of a set of 46 human ECD-hFc-8xHis proteins purified using the automated workflow. A. Workflow for protein quantification using the Bradford assay. B. Quantification of 46 purified proteins via Bradford assay using purified BSA to generate a standard curve. C. Coomassie-stained gel of protein samples (left) with purified BSA concentration standards (right). Green asterisk, protein 373 has a barely visible band at ∼90 kD. Blue asterisk, there is no band in the correct MW range for protein 565. Bands of the correct size could be visualized by Western blotting for proteins 373 and 565.

Concentration normalization of proteins for downstream applications is carried out using PBS as the diluent (Fig. 9A). A workbook is generated that maps the location and concentration of each stock protein in the rack of 96 screw cap tubes. Normalization scripts are executed on the Tecan Freedom Evo to calculate and dispense the appropriate volumes of protein and diluent into each well of a target working plate, ensuring uniform concentrations across all samples.

### Application of Automated Workflow to Generate Human ECD-hFc-8xHis Proteins

With an end-to-end workflow in place for generating assay-ready purified proteins, we applied it to produce 46 human ECD-hFc-8xHis proteins. Transfection of the 46 plasmids was performed in both the top and bottom half of 96-DWPs such that each plasmid was transfected in duplicate wells. Purified proteins were quantified using the Bradford assay (Fig. 9B). For our proteins, concentrations ranged up to 800 µg/ml, corresponding to a yield of 320 µg from a 4 ml culture. The Bradford assay is reliable for protein concentrations down to about 100 µg/ml. However, for proteins in our preparations with an apparent concentration of <200 µg/ml, the Bradford assay does not provide an accurate estimate of the concentration of the desired protein species.

There are always contaminants in the preparation, which run as multiple bands, and if each contaminant band is present in amounts similar to that of the desired protein band, the Bradford is then recording primarily the concentration of the contaminants (for example, protein 460 in Figure 9C). For such proteins, detection and quantification requires analysis by SDS-PAGE and densitometry.

In Figure 9C, we show SDS-PAGE analysis of the set of 46 test proteins from the upper half of the transfection plate. Proteins were visualized by Coomassie Blue staining. All but one of the proteins showed a band at the predicted size (although one band was barely visible). One protein had no detectable band in the correct molecular weight range. The protein with a barely visible band and the protein with no detectable band are denoted by asterisks in Figure 9C. These two ECD-hFc-8xHis proteins (which also contain a FLAG epitope tag) were visible on a Western blot via anti-FLAG staining (data not shown).

## Discussion

In this paper, we describe a modular automated system that is capable of end-to-end execution of a combined workflow that begins with transformation of plasmid DNA and ends with assay-ready purified desalted protein at the concentration required for downstream assays. We developed this platform to allow us to express and purify >2000 secreted human Fc fusion proteins from Expi293 cells. However, individual workflows described here could be adapted to other types of proteins and other cell lines, and most of the same modules could be used to purify cytoplasmic or nuclear proteins, or proteins bearing other affinity tags.

The system described here is composed of modular instruments that can be linked together in flexible arrays into a complete robotic platform. These components are inexpensive compared to commercially available single-application instruments that perform only one step in the overall workflow (e.g., plasmid purification). This makes our system particularly useful for basic research, since similar platforms could be assembled from units used for other types of experiments. Liquid-handling robots, incubators, and centrifuges are available in many academic facilities, or can be purchased as used equipment, and could be repurposed to build a modular automation platform for a relatively low cost.

To develop the workflow, we used mCherry-hFc-8xHis as a surrogate for ECD-hFc-8xHis proteins. Once the workflows were optimized, we used the platform to generate 46 purified ECD-hFc-8xHis proteins in duplicate including an mCherry-hFc-8xHis (i.e., 94 transfections, purifications and downstream processing steps). The 46 proteins were selected because similar versions (differing only in their C-terminal epitope tags) were produced for our previous cell-surface interactome screen (Wojtowicz et al. 2020).This screen was executed by researchers at Stanford and Caltech using a liquid-handling robot in the Caltech Protein Expression Facility (PEC) in the Beckman Institute, where the current system is also located. We created the 46-protein set for the purpose of developing and optimizing a new high-throughput multiplex methodology for screening for protein-protein interactions. This methodology, denoted as the MPIA, is described in a separate manuscript, posted to bioRxiv.

In this paper, we describe the in-house engineered robotic platform, automation workflows, and production of this set of 46 ECD-hFc-8xHis proteins, beginning with transformation of plasmid DNA. The platform successfully purified all 46 proteins, which cover a wide range of expression levels, ranging from barely detectable to ∼800 µg/ml.

### Modularity and Flexibility for Configuration and Workflow Diversity

The performance and utility of this automated platform are significantly enhanced by several key characteristics. The workflow is built around a central enclosure surrounded by two liquid handlers and various integrated modular automation devices, including a plate reader, centrifuge, plate sealer, plate peeler, decapper and recapper, as well as other devices not used in the workflow described here (e.g., plate washer). This modularity is a core design principle, allowing for flexibility in re-configuring different workflows as needed. This design avoids the need for acquiring highly specialized work cells dedicated to single tasks, as the integrated platform can accommodate a wide range of research and preclinical applications by reconfiguring its existing modules. Reconfiguration can be done rapidly, since each module is in its own BSL-2 cabinet on wheels and connects to the main BSL-2 central enclosure via an in-house designed custom magnetic docking system. The platform is engineered with modularity in mind, permitting individual devices to be detached for upgrades or repairs, thus supporting a future-proof and flexible system design. This characteristic also inherently allows for expandability to future workflows by adding new modules. Earlier concepts of such modular and flexible automation platforms have been proposed by others, and these ideas have influenced and driven lab automation technology to develop “plug-and-play” automation ready robot arm systems and user-friendly integration software (Wolf et al. 2024).

In future iterations of the workflows described in this paper, we will incorporate two additional pieces of equipment, which we have already procured. The first is an automated high speed shaking incubator (Kuhner MPS-Z) which has the capacity for 16 96-DWPs. The incubator is also housed in a modular BSL-2 dockable enclosure and will be fully integrated into the system. The second is a PF400 robotic arm, which has already been installed on our platform (Fig. 1), and can seamlessly access each device through the magnetic docking tunnels, facilitating loading, unloading, and transfer of plates between any docked devices. When this is done, it will be possible to conduct all of the workflows without manual transfer. We expect that this system will be capable of purifying 384 proteins (four 96-well plates) per week. We plan to integrate the platform with Laboratory Information Management Systems (LIMS) and expand its use to other complex biochemical and cell-based assays.

### Bio-Hazard Containment (BSL-2 Compliance)

A critical feature of the platform is its design to allow for bio-hazard containment in a BSL-2 environment. All modules are arranged to maintain operations within a BSL-2 sterile environment throughout the fully automated manipulation of samples, from DNA preparation to final protein purification. Specifically, all reagent handling, DNA-lipid complex formation, and distribution during mammalian cell transfection were performed in a sterile environment. Work with human HEK293-derived cell lines requires BSL-2 compliance. This ensures safety in general when working with human tissues, cells, or samples.

### User-Friendly Scheduling Software

All pipetting steps, incubation timings, and plate transfers for plasmid DNA preparation were carried out by the Hamilton Starlet liquid handler using the Hamilton Venus Instrument software 5. Additionally, normalization scripts were executed on the Tecan Freedom Evo to calculate and dispense appropriate volumes for DNA normalization. The “Green Button Go” scheduling software (GBG) from BioSero is a sophisticated but user-friendly automation software that allows seamless workflow automation between devices. We will also use this software to schedule the get and put movements of the PF400 SCARA robot arm.

### Benefits of Automation for High-Throughput Screens

The system’s automation enables a significant increase in the number of proteins produced in parallel, substantially reducing manual handling and processing time, and thereby accelerating the research timeline. The platform successfully transfected cells in 96-DWPs with plasmids encoding 46 ECD-Fc proteins and a fluorescent control protein (mCherry-hFc-8xHis) in duplicate, and purified and quantified the 94 proteins from the resulting supernatants in a single week.

The success of this automation approach will allow researchers to explore a wider range of experimental designs, leading to more thorough investigations when large protein libraries have to be produced in a timely manner to avoid protein expiration before assay execution. The entire process, from automated preparation of plasmid libraries (including DNA quantification and normalization) to efficient transient transfection of HEK293 cells, automated harvesting of cell culture supernatants, and subsequent affinity-based purification, is seamlessly integrated.

### Enhanced Reproducibility and Consistency

Another crucial advantage of this automated platform is its ability to improve reproducibility by minimizing human error and ensuring consistent processing conditions. The results confirm that the system achieved consistent DNA transfection grade plasmid DNA yields, accurate normalization, high transfection efficiency, and reproducible protein titers across multiple different proteins. DNA concentrations were consistently quantified using absorbance measurements on the Infinite 200 Pro plate reader, and normalization scripts on the Tecan Freedom Evo ensured uniform concentrations across all samples prior to downstream applications. Furthermore, plasmid DNA preparation was fully automated on the Hamilton Starlet liquid handling system, ensuring consistent execution of the plasmid purification protocol.

### Cost Savings through Miniaturization

The design of the automated platform, utilizing miniaturized volumes, leads to considerable cost savings on reagents and consumables. This efficiency is critical for high-throughput screening initiatives where reagent consumption can be substantial.

In conclusion, this study presents a modular, fully automated, 96-well plate-compatible workflow for protein production that minimizes human error and maximizes throughput, highlighting its flexibility, robustness, and BSL-2 compliance for diverse research and preclinical applications.

## Materials and Methods

### Plasmid transformation

Plasmid DNA transformation was partially carried out using manual steps, but full automation of this process is planned, using the robot platform. Competent DH5 alpha E. coli cells where prepared using the Mix & Go! E.coli Transformation Kit and Buffer Set, following the Manufacturer’s instructions (Zymo Research, Cat.# T3002). Best transformation results were obtained by preparing the E. coli cells and freezing them in bulk volumes of 5 ml aliquots at −80 °C. Immediately before the transformation cells were thawed at RT and dispensed into 96-well PCR plates using the liquid handler or multichannel pipette if done manually, adding 20 ul to each well. DH5 alpha competent E. coli cells were transformed by adding 2 ul of plasmid DNA to the cells, followed by adding 100 ul SOC or LB medium and mixing gently. After a 1 hour recovery incubation period at 37°C transformed cells were plated using an automated Tecan Evo-based spreading method to ensure uniform colony distribution across agar plates. Individual colonies were then hand-picked and inoculated into selective liquid media for overnight growth. These cultures were stored as glycerol stocks and stored at –80 °C as a renewable source. When needed, the glycerol stocks were revived to inoculate fresh deep-well 96-well plates to renew the glycerol stocks when running low.

### Generation of transfection-quality plasmid DNA

Plasmid DNA preparation was performed using the Macherey-Nagel NucleoBond Xtra 96-well midi prep kit. The protocol was fully automated on the Hamilton Starlet liquid handling system. Overnight bacterial cultures (1ml per well) were grown in four replicate 96-DWPs. Bacteria were pelleted by centrifugation, resuspended in RES buffer, and the four replicate plates were pooled into a single 96-DWP plate. Cell lysis was induced by the addition of LYS buffer followed by a brief incubation. The lysed cell suspension was then neutralized using NEU buffer, and the lysate was clarified first by spinning in the robot centrifuge at 1000 x g twice and then by vacuum filtration through the lysate filtration plate (under these conditions up to 4 deep well plates of culture could be accommodated with one single lysate filtration plate). The cleared lysate was then transferred to the pre-equilibrated NucleoBond® Xtra EF Plate and the plasmid DNA captured allowing gravity flow of the lysate through the plate. The bound DNA was then washed 2X with buffer Buffer ENDO-EF followed by one wash with buffer WASH-EF under gravity flow. Plasmid DNA was then eluted into a 96-DWP adding buffer ELU-EF under gravity flow. The eluted plasmid DNA was then precipitated by adding 100% isopropanol and transferred to the equilibrated prewashed Finalizer Plate spin plate. The plasmid DNA-precipitate retained in the Finalizer Plate was washed with 80% Ethanol using the vacuum manifold followed by a 1000 x g spin. The precipitate was allowed to redissolve for 5 minutes by adding with sterile water provided by the kit and preheated to 70°C and was retrieved by spinning the Finalizer Plate at 1000 x g in the robot plate centrifuge into a PCR 96-well plate.

Following the plasmid prep, DNA concentrations were quantified using A260/280 absorbance measurements on UV-Star assay plates (Greiner Bio-One) read by the Infinite 200 Pro plate reader. A spreadsheet was then generated containing all DNA concentrations, which was subsequently used as input for a normalization workbook. This workbook calculated the precise volumes of DNA and diluent (Opti-MEM) required to normalize all samples to a final concentration of 20 ng/µL for later transfection into Expi293 cells.

### DNA Quantification and Normalization

Plasmid DNA was quantified using a UV-Star® 384-well UV-transparent microplate (Greiner Bio-One), designed for high-throughput UV absorbance measurements. Purified plasmid DNA samples were diluted 1:10 in 1× TE buffer (pH 7.4) by mixing 2 µL of DNA with 18 µL of buffer and loaded in triplicate into quadrants 1, 2, and 3 of the plate. A standard curve was generated using mCherry plasmid DNA at concentrations of 200 ng/µL, 50 ng/µL, 12.5 ng/µL, and 3.125 ng/µL, each diluted in TE buffer. For each standard, 20 µL of the diluted DNA was added directly to the wells. Absorbance was measured at 260 nm for DNA quantification, and at 280 nm and 230 nm to assess protein and organic contamination, respectively. DNA concentrations were calculated using the Beer–Lambert law, applying a conversion factor of 50 ng/µL per A260 unit for double-stranded DNA. All absorbance measurements were performed using a UV-Vis microplate reader with pathlength correction enabled where applicable, and care was taken to avoid air bubbles and ensure consistent pipetting across replicates.

### Expi293 Cell Transfection

Expi293 cells were cultured in Expi293 Expression Medium (Thermo Fisher Scientific) under standard conditions: 37°C, 8% CO₂, and >80% humidity. Cells were maintained at a viability greater than 98% prior to transfection. For transfection, cells were seeded into 96-well plates at a density of 4 × 10^5^ cells/mL. Transfections were carried out using the ExpiFectamine 293 Transfection Kit (Thermo Fisher Scientific). Plasmid DNA was added at 1 µg per mL of expression volume. DNA-lipid complexes were prepared by diluting the plasmid DNA and ExpiFectamine reagent in Opti-MEM, followed by a 3-minute incubation. After complex formation (5 minutes), the mixtures were automatically distributed to the cell plates using the Hamilton Starlet. Transfected plates were then incubated at 37 °C with 8% CO₂ and shaken at 1000 rpm on a 3 mm radius orbital platform shaker. Enhancers provided in the kit were added 17–20 hours post-transfection according to the recommended protocol. All reagent handling, complex formation, and distribution were performed in a sterile environment to maintain BSL-2 compliance.

Following transfection, cells were incubated in a humidified environment without additional media exchange. Expression proceeded for 4.5 days, during which plates were continuously shaken at 1,000 RPM on an orbital shaker with a 3 mm radius. Plates remained undisturbed inside the incubator to minimize variation due to handling. At the end of the expression period, the supernatant was harvested by spinning at 1000 x g using the robotic plate centrifuge and directly transferred from the 96-well plates using the Hamilton Microlab Starlet, equipped with tip-based aspiration to avoid disturbing the settled cells. Clarified supernatant was stored in fresh deep-well plates for downstream purification.

### Protein Purification

Cell supernatant was harvested 4.5 days post transfection and filtered through a 96 Well PALL 1 ml, 1.2um pore size polyethersulfone (PES) based filter plate. Filter plates were subjected to centrifugation at 1000g for 1 minute. Protein purification was performed using a Ni-NTA metal chelate affinity resin on the Hamilton. For the purification, 20 ml of resin bed volume was prepared and equilibrated with 1 x PBS. Clarified supernatants (1 ml per well) were incubated with 50 µL of equilibrated resin in 96W deep-well plates. Incubation was carried out at room temperature for 1 hour with intermittent resin resuspension every 5 minutes by automated agitation to facilitate protein binding to the Ni-NTA resin. The resin and supernatant mixture was then transferred in 500 µL increments to a 96-well Pall filter plate with a 1.2 µm pore size PES membrane, where resin packed to form an affinity column bed using 3 minute 300 x g spins in the robot plate centrifuge. The resin was washed three times with 500uL of 1x PBS using 1000 x g to remove nonspecifically bound proteins. His-tagged proteins were eluted with two 200 µL elution buffer volumes containing 500 mM imidazole and collected in a clean 96-well plate by spinning at 1000 x g for 3 minutes.

### Desalting of Protein Samples

Plasmid DNA samples were desalted using the Zeba™ 96-Well Desalting Plate, 40K MWCO, 0.1 mL (ThermoFisher Scientific, Cat. No. 89810), which utilizes size-exclusion chromatography (SEC) to efficiently remove low-molecular-weight contaminants such as salts, enzymes, and small buffer components while retaining plasmid DNA. Each well was first equilibrated with 250 µl of 1× PBS buffer (pH 7.4) to remove the storage solution and condition the resin. Equilibration was performed by centrifuging the plate at 700 × g for 2 minutes. This washing step was repeated three times to ensure complete removal of preservatives. After equilibration, 100 uL of protein sample was carefully loaded onto the center of the resin bed in each well. The plate was then centrifuged at 700 × g for 3 minutes to elute the desalted protein sample, which was collected into a clean receiver plate placed beneath the column plate. All steps were performed at room temperature, and care was taken to avoid disturbing the resin during sample application and centrifugation.

### Protein Quantification and Normalization

Protein samples were prepared starting with spinning the rack of 96 screw cap tubes of stock protein, followed by decapping the rack using a De/Recapper. 1x PBS was then dispensed into all wells of a 384-well plate using a liquid handler. Next, the desalted stock protein was dispensed into quadrant well 1 of the 384-well plate. Three serial dilutions were performed across quadrant wells 2 to 4 to achieve a range of protein concentrations. The 96 screw cap tubes containing the stock protein were recapped with the De/Recapper after dispensing. Protein Assay Dye Reagent Concentrate (Bio-Rad, Cat. No. 5000006) was added to all wells of the 384-well plate, and the plate was read at 595 nm to determine protein concentrations. Using these concentrations, appropriate volumes of PBS and protein were dispensed into a 96-well PCR plate to normalize protein amounts across samples. The 96 screw cap tubes of stock protein were recapped again after dispensing the protein volume. Finally, the 96-well PCR plate containing the normalized protein samples was sealed in preparation for further analysis.

### SDS-PAGE

SDS-PAGE was performed to assess the purity and integrity of the normalized protein samples. Equal volumes of normalized protein samples from the sealed 96-well PCR plate were mixed with 4× Laemmli sample buffer containing 100 mM dithiothreitol (DTT) and heated at 95 °C for 5 minutes to denature the proteins. Samples were then loaded onto 4–20% gradient polyacrylamide gels and separated at a constant voltage of 200V for approximately 45 minutes in 1× Tris-Glycine-SDS running buffer. Following electrophoresis, gels were stained with Coomassie Brilliant Blue R-250 for visualization of total protein.

## Acknowledgments

This work was funded by an NIH TRO1 grant, GM150125, to K.Z., K. Christopher Garcia (Stanford), and Matthew Thomson (Caltech). Maxine Wang was supported by a fellowship from the H.S. Chau Foundation. Work at the Caltech Protein Expression Center is also supported by the Beckman Institute at Caltech. Some of the equipment was purchased with grants to the Merkin Institute and Chen Foundation at Caltech, and from funds from the BBE Division and the Provost’s Office.

Conceptualization, J.V., W.M.W., P.V., M.A.A., K.Z.; Investigation, P.V., J.V., M.A.A., A.W.L., A.Z., E.G., M.L.W.; Data Curation and Visualization: J.V., W.M.W., P.V.; Writing and Editing, J.V., W.M.W., K.Z.; Supervision, J.V., K.Z., W.M.W.

We thank Matt Thomson, Chris Garcia, Pamela Bjorkman, Barbara Wold, and Viviana Gradinaru for helpful discussions.

